# *ALMS1* KO rat: a new model of metabolic syndrome with spontaneous hypertension

**DOI:** 10.1101/2024.09.22.614364

**Authors:** Ankita B. Jaykumar, Sumit R. Monu, Mariela Mendez, Nour-Eddine Rhaleb, Pablo A. Ortiz

## Abstract

ALMS1 is a protein initially associated with Alström syndrome. This is a rare human disorder characterized by metabolic dysfunction, hypertension, obesity and hyperinsulinemia. In addition, *ALMS1* gene was linked to hypertension status in a multipoint linkage population analysis. However, the mechanisms by which ALMS1 contributes to the development of obesity, insulin resistance and other metabolic disturbances are unknown. To study the role of ALMS1 in blood pressure regulation and renal function we previously generated an ALMS1 knockout rat model, where we found these rats are hypertensive. In this study, we further characterized the *ALMS1* knockout rat, and found that they exhibit most characteristics of metabolic syndrome including hypertension and higher body weight by 10-12 weeks of age. In contrast, obesity, hyperinsulinemia and vascular dysfunction manifested at around 14-16 weeks of age. Interestingly, *ALMS1* KO rats developed hyperleptinemia prior to the development of obesity rapidly after weaning by 7 weeks of age, suggesting an early role for ALMS1 in the hormonal control of leptin. We also found that female *ALMS1* KO rats develop severe metabolic syndrome with hypertension similar to their male counterparts, lacking any protection often associated with better cardiovascular outcomes. Therefore, ALMS1 is an essential gene for sex-and age-dependent metabolic function. The *ALMS1* knockout rat provides an invaluable pre-clinical animal model that recapitulates most symptoms present in patients and allows the study of new drugs and mechanisms that cause metabolic syndrome.

## Introduction

Metabolic syndrome is a clustering of cardiovascular risk factors associated with hypertension, insulin resistance, glucose intolerance, obesity, hypertriglyceridemia, hypercholesteremia and low levels of high-density lipoprotein. Pre-menopausal women are relatively protected from metabolic syndrome compared to men. In contrast, postmenopausal women have a higher prevalence of metabolic dysfunction, with the age-related loss of female sex hormones [1-3]. Several genetic loci and genes have been discovered that are individually associated with one or a few of the above-mentioned phenotypes of metabolic syndrome. To our knowledge, there is no single gene that, when deleted in animals causes most phenotypes associated with MetS, including hypertension.

Alström syndrome is a rare, autosomal recessive genetic disorder linked to deleterious mutations in the *ALMS1* gene, which causes multiple phenotypes that are risk factors for metabolic syndrome in addition to cardiovascular disease, and chronic kidney disease. In Alström syndrome patients renal function deteriorates progressively with age and end-stage renal disease is a common cause of death [4-7]. Hypertension is a common risk factor for chronic kidney disease (CKD) [8,9]. SNPs in the ALMS1 gene are linked to decreased renal function (CKD gen) and were also linked to hypertension and increased pulse pressure status in a multipoint linkage analysis in 7 primary sibling samples of African American, Caucasian and Mexican populations [10]. This suggests that ALMS1 is a gene with deep penetrance on a wide cluster of metabolic phenotypes and could be an essential player in maintaining normal metabolic health.

We previously showed that ALMS1 is involved in blood pressure regulation and control of renal function by promoting endocytosis of the Na^+^/K^+^/2Cl^-^ cotransporter NKCC2 in the kidney [11]. Another study showed that *ALMS1* mutant mice displayed altered intracellular localization of glucose transporter-4 (GLUT4) and decreased insulin-stimulated trafficking of GLUT4 to the plasma membrane in adipocytes [12], suggesting a mechanism for the development of insulin resistance due to *ALMS1* deletion. In addition, ALMS1 KO mice develop early onset obesity and insulin resistance [13]. However, it is unclear whether ALMS1 deletion causes the whole range of symptoms of metabolic syndrome including obesity, insulin resistance, hyperleptinemia, and altered lipid profile, in addition to hypertension.

In this study, we characterized an *ALMS1* knockout rat, generated on the genetic background of the Dahl Salt sensitive rat. Our data show that this model exhibits age-dependent and sex-independent obesity, hypertension, hyperleptinemia, hyperlipidemia and vascular dysfunction. Therefore, this *ALMS1* knockout rat spontaneously and without any dietary intervention, exhibits all phenotypes of metabolic syndrome making it a very valuable tool for studying the mechanisms and new drugs to treat metabolic syndrome. Our data also point to the pleiotropic role of ALMS1 in metabolism and vascular function.

## Result

### Generation of *ALMS1* genetic deletion rat model

We studied the effect of *ALMS1* deletion in rats on metabolic parameters, this transgenic rat was generated using zinc finger nuclease gene editing technology in collaboration with the Gene Editing Rat Resource Center (GERRC) at the Medical College of Wisconsin [11]. Deletion of 17 base pairs in exon 1 of the *ALMS1* gene results in a premature stop codon to generate the *ALMS1* deletion in Dahl salt sensitive rats (**Figure 1A**). **Figure 1B** shows a 90% reduction in ALMS1 expression in kidney lysates from *ALMS1* KO rats. This indicates successful whole-animal knockout of ALMS1 supporting the use of these rats for studying the effect of *ALMS1* deletion on metabolic phenotype.

**Figure 1:**
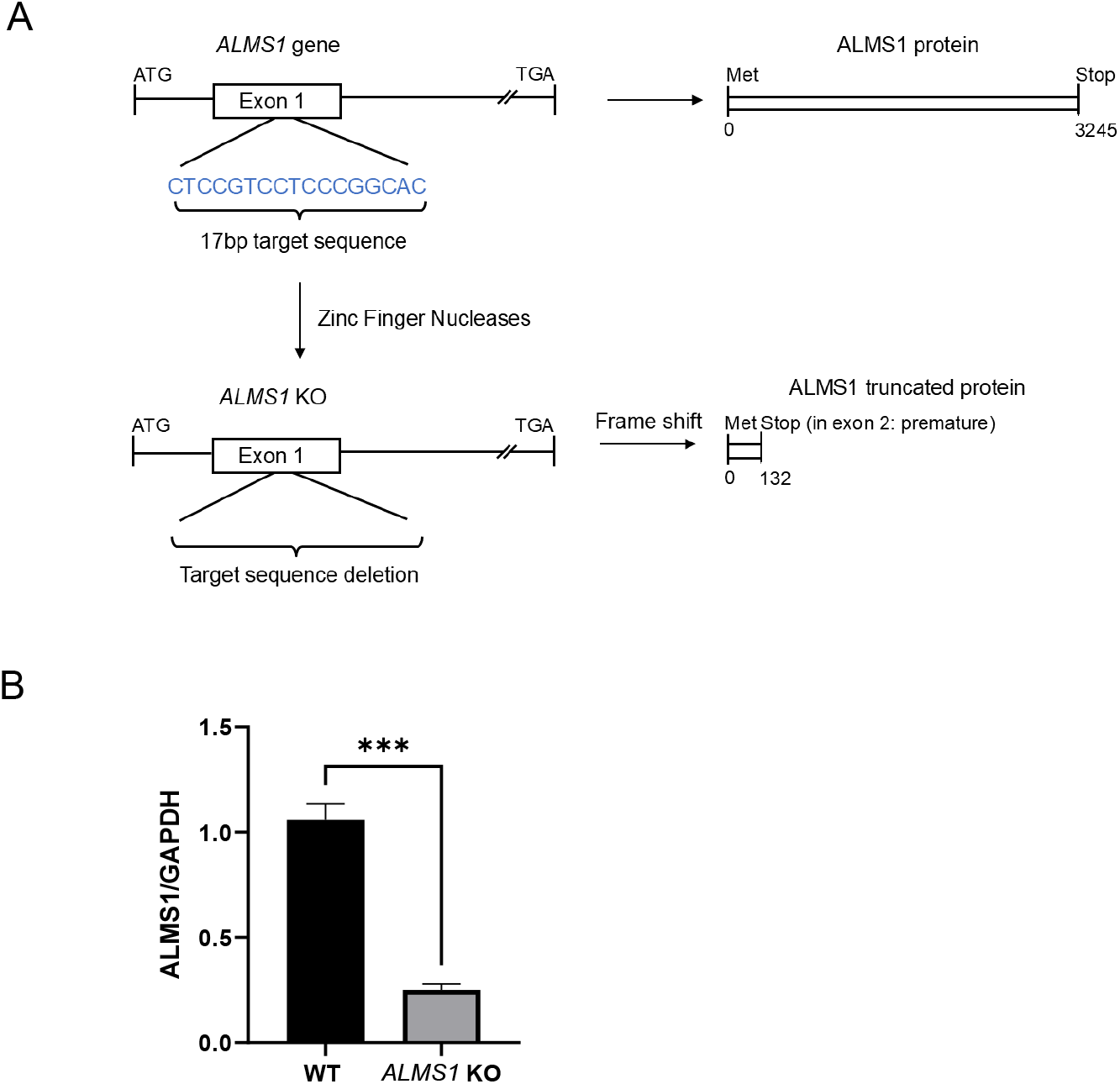
Generation of *ALMS1* KO rats. **A**) Zinc finger nucleases target 17 base pairs in exon 1 of the ALMS1 gene leading to a frameshift and causing a pre-mature stop codon in exon 2 (figure adapted from [11]). **B**) Representative Western blot for ALMS1 in kidney lysates from WT and *ALMS1* KO rats, n = 3, * p < 0.05 vs. WT. Data is represented as mean ±SEM; analyzed by a two-tailed Student’s *t*-test.

### *Young ALMS1* KO rats (7-9 weeks old) do not develop metabolic syndrome

Previous reports have shown that transgenic *ALMS1* mice are characterized by age-dependent obesity, metabolic syndrome and kidney damage [13]. We measured the body weight of wild-type (WT) and *ALMS1* KO rats and Table 1 clearly shows that at age 7 weeks, body weight was not different between the sexes or the strains. We found no difference in plasma insulin or blood glucose levels between the strains or the sexes. Interestingly, plasma leptin was extremely higher at 6–7 weeks of age in male rats (ALMS1: 3530 ± 300 vs. WT: 172 ± 16 pg/ml) as well as in female rats (ALMS1: 2241 ± 523 vs. WT: 76 ± 24 pg/ml) (**Table 1**). These data indicate that at 6-7 weeks of age, *ALMS1* KO rats do not significantly develop metabolic syndrome. However, elevated plasma leptin in young *ALMS1* KO rats occurs before developing metabolic dysfunction. This points to *ALMS1* as a potentially novel gene involved in leptin biology/function, expression and/or release.

**Table 1:** 6-7-weeks old ALMS1 KO rats have a normal metabolic profile. Blood glucose, plasma insulin, and body weight in WT and ALMS1 KO rats measured in 6-7-weeks of age were similar except for the plasma leptin which was elevated in the *ALMS1* KO rats. All values represent mean ± SEM and statistical analysis was performed with two-tailed Student’s *t*-test. Male WT n=9; male *ALMS1* KO n=6; Female WT n=8; Female *ALMS1* KO n=7; *p<0.05 vs. WT.

| Metabolic parameters | WT- male (6-7-weeks old) | ALMS1 KO- male (6-7-weeks old) | WT- female (6-7-weeks old) | ALMS1 KO-female (6-7-weeks old) |
| --- | --- | --- | --- | --- |
| Body weight (g) | 202±7 | 215±6 | 150±5 | 165±7 |
| Plasma leptin (pg/ml) | 172±16 | 3530±300 * | 76±24 | 2241±523* |
| Plasma insulin (mg/ml) | 2±0.4 | 3±1 | 1.9±0.3 | 3.4±0.7 |
| Blood glucose (mg/dl) | 98±5 | 116±9 | 93±5 | 98±4 |

### 14-16 weeks old *ALMS1* KO rats develop hyperphagia and obesity

Since 6–7-weeks-old *ALMS1* KO rats did not have a higher body weight than wild-type littermates, we tested whether older *ALMS1* KO rats were obese. We measured their body weights and found that both male and female *ALMS1* KO rats have a significantly higher body weight than their wild-type littermates, starting at age week 9 in males and week 7 in females (**Figure 2A**). In 16-18-weeks-old ALMS1 rats, we measured abdominal fat mass and found that it was higher in both male (*ALMS1* KO: 31.3 ± 3.4 g vs. WT: 11.36 ± 2.5 g) and female *ALMS1* KO rats (*ALMS1* KO: 35.8 ± 3.7 g vs. WT: 7.62 ± 1.8 g, n = 3) compared to their wild type littermates (**Figure 2C, 2D**) and appeared obese at 16 weeks of age (representative image of female rats; **Figure 2B**). We then measured food intake in 16–18-weeks-old *ALMS1* KO rats and found that both male (*ALMS1* KO: 27 ± 3 g/day vs. WT: 19 ± 2 g/day) and female *ALMS1* KO rats (*ALMS1* KO: 25 ± 2.5 g/day vs. WT: 13 ± 1 g/day) are hyperphagic (**Figure 2E, 2F**), a potential cause for their obesity. We then measured plasma leptin levels in 16-18-weeks-old *ALMS1* KO and found that both male (male *ALMS1* KO: 68300 ± 10500 vs. WT: 2750 ± 1100 pg/ml) and female *ALMS1* KO rats were hyperleptinemic (female *ALMS1* KO: 64100 ± 15900 vs. WT: 900 ± 300 pg/ml) (**Figure 2G, 2H**). These data suggest that *ALMS1* KO rats may have a progressive deficiency in leptin signaling leading to blunted satiety, suggestive of leptin resistance.

**Figure 2:**
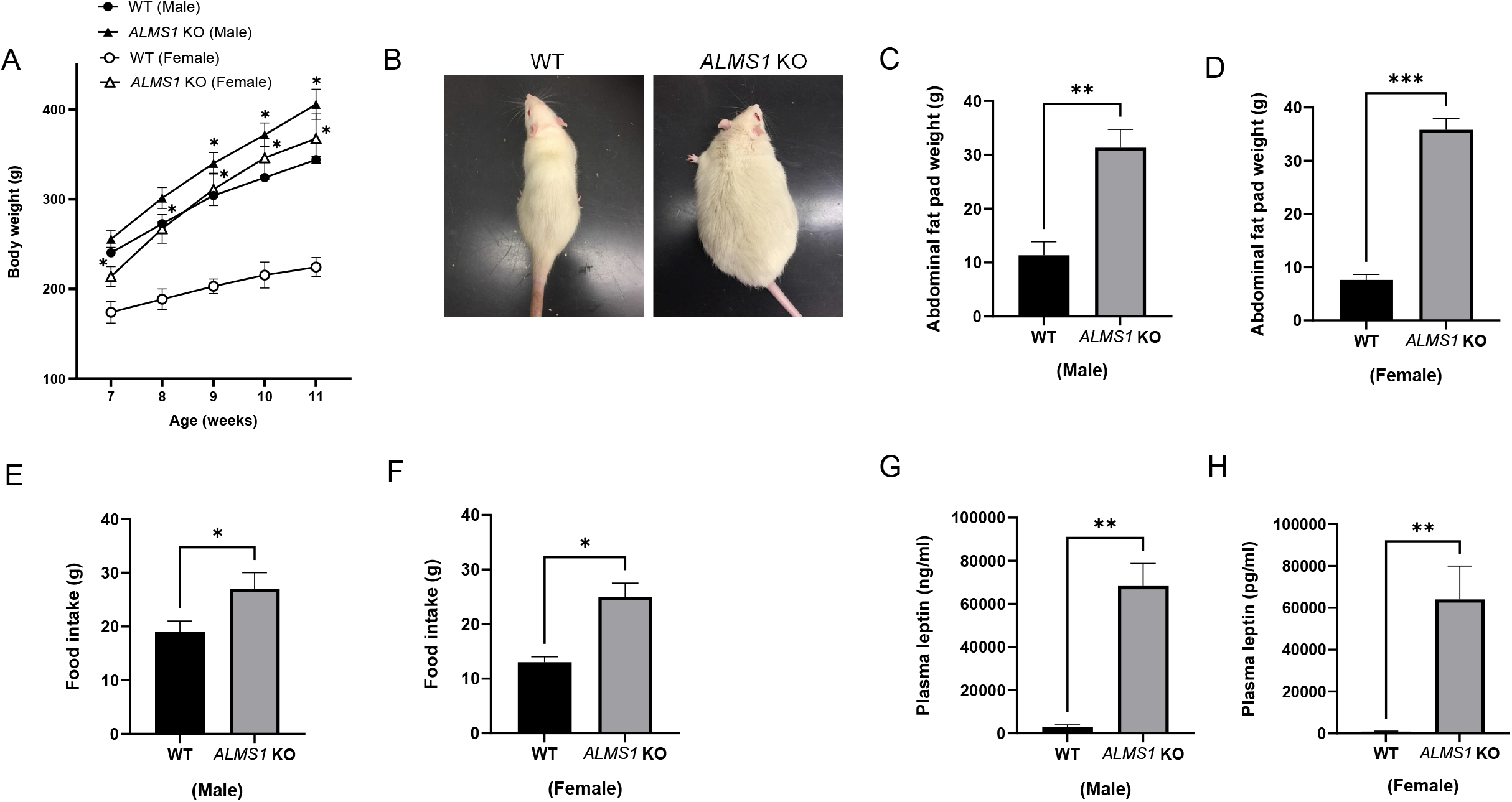
14-16 weeks old *ALMS1* KO rats develop hyperphagia and obesity. **A**) Body weight differences between the sexes of *ALMS1* KO rats, n = 5 vs. WT. **B**) Representative image of old female *ALMS1* KO and WT rats. **C&D**) Abdominal fat pad weight in both sexes of *ALMS1* KO rats, n = 3 vs. WT. **E&F**) Daily average food intake in both sexes of *ALMS1* KO rats, n = 5 vs. WT. **G&H**) Plasma leptin concentration in both sexes of *ALMS1* KO rats, n = 4 vs. WT. Data is represented as mean ±SEM; analyzed by one or two-tailed Student’s *t*-test; *p<0.05, **p<0.01 ***p<0.0005.

### *ALMS1* KO rats exhibit altered lipid profile

Metabolic syndrome is associated with several abnormalities in blood lipid profile such as serum triglycerides, low-density lipoprotein (LDL), high-density lipoprotein (HDL), and total cholesterol [4-7]. We observed that serum from 16-18 weeks old *ALMS1* KO rats appeared milky, suggesting hyperlipidemia (**Figure 3A**). Therefore, we measured their serum lipid profile and found that triglyceride levels were significantly elevated in both male (*ALMS1* KO: 543.7 ± 179.3 mg/dl *vs*. WT:100.3 ± 15.5 mg/dl) and female *ALMS1* KO rats (*ALMS1* KO: 595.9 ± 200.1 mg/dl *vs*. WT: 55.6 ± 12.5 mg/dl). Significant differences were observed in serum total cholesterol (*ALMS1* KO: 166 ± 41.5 mg/dl vs. WT: 81.9 ± 4.9 mg/dl) HDL (*ALMS1* KO: 41.9 ± 7.9 mg/dl vs. WT: 67.8 ± 2.2 mg/dl) LDL (*ALMS1* KO: 49.5 ± 17.6 mg/dl vs. WT: 15 ± 1.4 mg/dl) only in female *ALMS1* KO rats, but not in male *ALMS1* KO rats (**Figure 3B, 3C**). These data indicate that ALMS1 KO rats develop progressive hyperlipidemia with several lipid profile parameters such as serum HDL, LDL and total cholesterol levels particularly exacerbated only in females but not male *ALMS1* KO rats. These data suggest that ALMS1 may be involved in sex-based differences in lipid metabolism.

**Figure 3:**
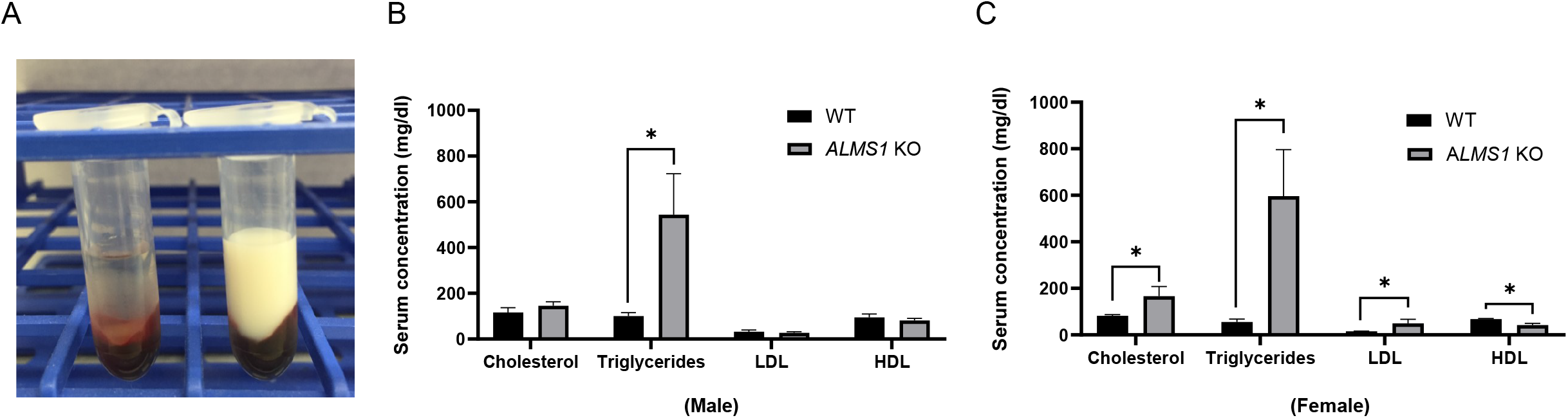
*ALMS1* KO rats exhibit altered lipid profiles. **A**) Males *ALMS1* KO: n = 7, WT: n = 4, * p < 0.01 vs. WT and **B**) Females *ALMS1* KO: n = 7, WT: n = 4, * p < 0.05 vs. WT. Data is represented as mean ±SEM; analyzed by a one-tailed Student’s *t*-test.

### *ALMS1* KO rats are insulin resistant

Insulin resistance is characterized by an impaired insulin sensitivity due to decreased ability of cells to respond to insulin and regulate blood glucose levels [1-3]. We tested whether depletion of *ALMS1* in rats reproduced insulin resistance as seen in the *ALMS1* mutant mice model [13]. We measured random blood glucose by tail snips, and we found that it was not different between the groups in either male (*ALMS1* KO: 134.5 ± 9.4 mg/dl vs. WT: 132.3 ± 9.9 mg/dl) or female rats (*ALMS1* KO: 122 ± 2.5 mg/dl vs. WT: 130.3 ± 2.4 mg/dl) (**Figure 4A, 4B**). However, the fasting blood glucose measurement showed significantly higher values in both male (*ALMS1* KO: 108 ± 4 mg/dl vs. WT: 61 ± 17 mg/dl) and female rats (*ALMS1* KO: 100 ± 12 mg/dl vs. WT: 71 ± 6 mg/dl) rat groups (**Figure 4C, 4D**). Fasting plasma insulin was measured to be significantly higher in both female and male KO rats (Male-*ALMS1* KO: 13.2 ± 5.6 ng/ml vs. WT: 1.095 ± 0.6 ng/ml) and (Female-*ALMS1* KO: 7.6 ± 1.7 ng/ml vs. WT: 0.82 ± 0.27 ng/ml) (**Figure 4E, 4F**), indicating that *ALMS1* KO rats are hyperinsulinemic. Together, these data suggest that *ALMS1* KO rats may have deficient insulin signaling, leading to their decreased ability to lower fasting blood glucose, indicative of insulin resistance.

**Figure 4:**
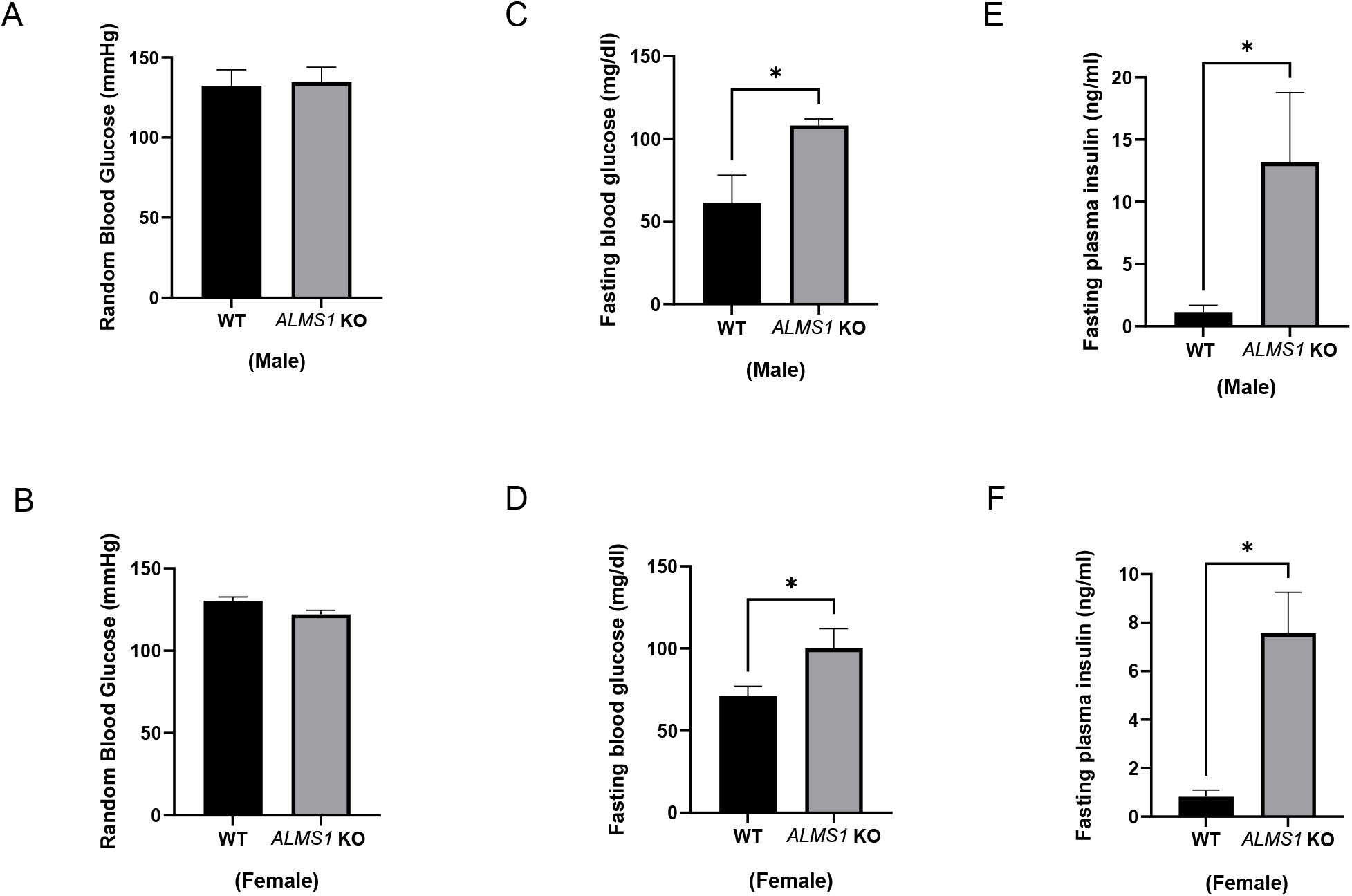
*ALMS1* KO rats are insulin resistant. **A&B**) Random blood glucose in both the sexes of *ALMS1* KO rats, n = 3, **C&D**) Fasted-blood glucose in both sexes of *ALMS1* KO rats, n = 3, * p < 0.05 and **E&F**) Plasma insulin in both sexes, male *ALMS1* KO: n = 5 & WT: n = 3, female *ALMS1* KO: n = 9 & WT: n = 6, * p < 0.05 vs. WT. Data is represented as mean ±SEM; analyzed by one or two-tailed Student’s *t*-test.

### Male and female ALMS1 KO rats are hypertensive

Hypertension is a significant risk factor in determining the severity of metabolic syndrome [14-19]. To test whether genetic deletion of *ALMS1* in rats leads to a sex-dependent increase in blood pressure, we used 7-9-weeks old *ALMS1* KO rats and WT littermates fed on a regular salt diet (0.22% Na) to measure systolic blood pressure by tail-cuff plethysmography. We found that both male (*ALMS1* KO: 151 ± 5.1 mmHg vs. WT: 125 ± 4.2 mmHg) and female (*ALMS1* KO: 199 ± 7.1 mmHg vs. WT: 134 ± 3.3 mmHg) *ALMS1* KO rats are hypertensive (**Figure 5A**). Systolic blood pressure in female *ALMS1* KO rats was higher than that of their male counterparts (**Figure 5B**). This suggests that blood pressure is more severely elevated in female ALMS1 KO rats than males. This finding in ALMS1 KO rats on a Dahl SS background is interesting because, previous studies have shown that female Dahl SS rats are protected from hypertension in comparison to male Dahl SS rats [20]. This indicates that deletion of ALMS1 eliminates any protection of hypertension that was observed in female rats.

**Figure 5:**
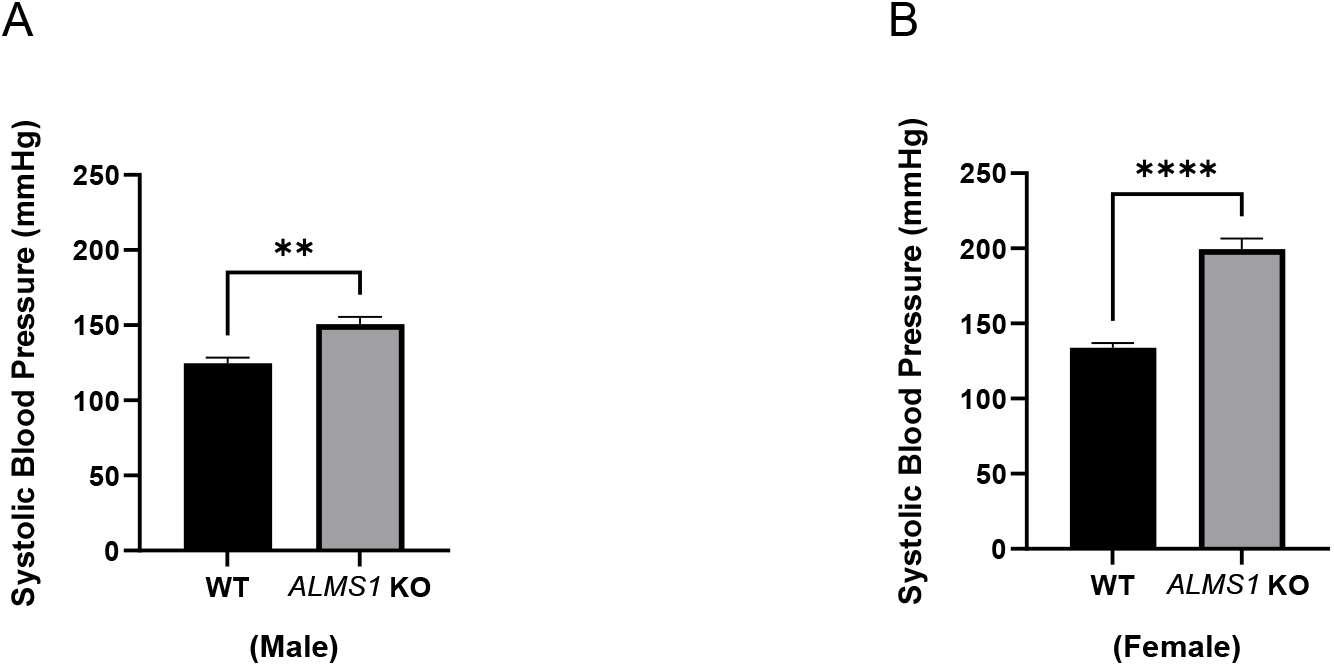
*ALMS1* KO rats are hypertensive. **A&B**) Systolic blood pressure in both sexes of *ALMS1* KO rats; Females-n = 6, **** p < 0.0001 vs. WT. B) Males-WT: n = 5 and ALMS1 KO: n=8, **p < 0.005 vs. WT. Data is represented as mean ±SEM; analyzed by two-tailed Student’s *t*-test.

### *ALMS1* KO rats exhibit vascular dysfunction

The role of ALMS1 in vascular reactivity is not known. Each component of the metabolic syndrome has been shown to affect the vascular endothelium and smooth muscle cell function to cause vascular dysfunction and disrupt vascular homeostasis [21]. Given the metabolic pathophysiology in *ALMS1* KO rats, we asked whether the deletion of *ALMS1* in rats led to vascular dysfunction. We measured the vasorelaxation response of thoracic aortic rings prepared from WT and *ALMS1* KO rats to methacholine- and nitroprusside. This is a measure of endothelial-dependent and endothelial-independent vasorelaxation, respectively [22]. Given that older rats develop metabolic dysfunction, we tested the vascular reactivity of rats between the ages of 14-20 weeks with significant metabolic defects. We found that thoracic aorta from both 14-20 weeks-old female and male *ALMS1* KO rats had a reduced vascular reactivity to nitroprusside (**Figure 6A, 6B**). This indicated a defect in the relaxation response of arterial smooth muscle to nitric oxide donor in the aorta from *ALMS1* KO rats between the ages of 14-20 weeks old. However, it should be noted that male *ALMS1* KO rats showed a defect in nitroprusside-induced vasorelaxation at doses much smaller than that observed in female *ALMS1* KO rats. In addition, abdominal aortic rings from both female and male *ALMS1* KO rats showed a decreased endothelial-dependent vasorelaxation in response to methacholine (**Figure 6C, 6D**). Overall, these data together suggest that ALMS1 function regulates endothelial function in the context of metabolic dysfunction.

**Figure 6:**
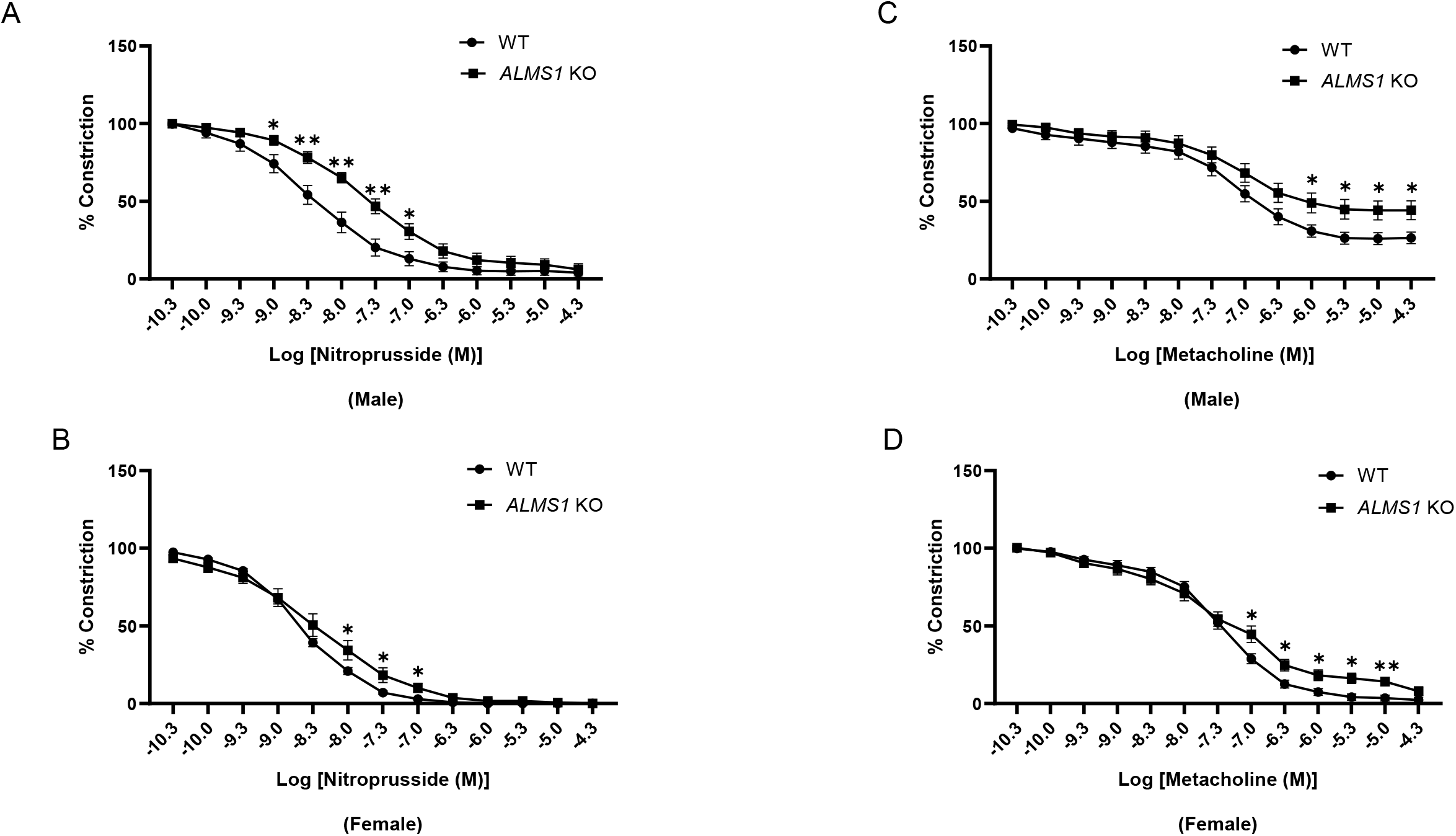
*ALMS1* KO rats exhibit vascular dysfunction. **A&B**) Nitroprusside-induced vasorelaxation in abdominal aorta from 14-20 weeks-old male and female *ALMS1* KO rats; n=6, *p<0.05, **p<0.005. **C&D**) Methacholine-induced vasorelaxation in abdominal aorta from 14-20 weeks-old male and female *ALMS1* KO rats; n=6, *p<0.05, **p<0.005. Data is represented as mean ±SEM; analyzed by student’s *t*-test.

## Discussion

The coexistence of hypertension and diabetes mellitus triples the risk for cardiovascular disease, as indicated in the Framingham Heart study [16] and MRFIT trials [17]. Obesity-related hypertension is also known to predispose to coronary heart disease, renal dysfunction, left ventricular hypertrophy and congestive heart failure [23-26]. The presence of hypertension in the setting of obesity and diabetes creates a vicious feed-forward loop that also increases the occurrence of end-stage kidney failure [27]. Despite the medical importance of metabolic syndrome, there are very few animal models that recapitulate all hallmarks of this syndrome, including hypertension. In this manuscript we present data showing that deletion of ALMS1 in rats, causes all phenotypes of metabolic syndrome, while on a normal diet, including hypertension, independently of sex. There is significant evidence that the ALMS1 gene is associated with metabolic diseases, hypertension and kidney disease in the general population. Patients with absence of functional ALMS1 exhibit exacerbated symptoms of metabolic syndrome including progression of cardiac and kidney disease [4-7]. Two independent genome-wide association studies showed single nucleotide polymorphisms in the *ALMS1* locus associated with lower GFR and chronic kidney disease [28-31]. Quantitative trait loci (QTL) associated with blood pressure and salt-sensitive hypertension were mapped to the *ALMS1* locus [32-36]. Thus, ALMS1 is likely to control multiple pathways involved in metabolism and cardiovascular function, and our data support this view.

Our study shows that *ALMS1* KO rats are an accurate model of metabolic syndrome, which includes hypertension and dyslipidemia. We show that 14-20 weeks-old male and female *ALMS1* KO rats are hyperinsulinemic, obese, hyperphagic, hypertriglyceridemic and hyperleptinemic. Female *ALMS1* KO rats also exhibit metabolic disturbances including higher serum cholesterol, LDL, and low HDL levels. Insulin resistance in the ALMS1 KO rat may develop secondarily to obesity, since we previously showed that young non-obese *ALMS1* KO rats are not hyperinsulinemic [11]. However, hyperleptinemia is not likely to develop secondary to obesity because 7 weeks old *ALMS1* KO rats have elevated plasma leptin levels before they show any signs of obesity. This suggests a primary role of ALMS1 in leptin production or signaling that causes hyperleptinemia. In addition, our data also support the hypothesis that hyperleptinemia and hence, leptin resistance could be the cause of obesity in our rat model. Previous studies have shown that in hypothalamic neurons, ALMS1 is known to be present in the base of the cilia, which plays a role in the regulation of satiety. In foz/ foz mice, loss of ALMS1 from this location coincided with a reduction in adenylyl cyclase 3 (AC3), a signaling protein implicated in obesity. In agreement with our data, another study showed that both sexes of global and mesenchymal stem cell-specific knockout of ALMS1 showed increased serum leptin concentrations [37]. This, together with our data suggests a role for ALMS1 in neuronal ciliary function and control of satiety [38]. Overall, our study suggests that the *ALMS1* KO rat provides an excellent new model to elucidate the associated pathophysiological mechanisms of metabolic dysfunction and cardiovascular risk factors.

ALMS1 is involved in ciliary function in some, but not all, cells. A study demonstrated that the knockdown of ALMS1 in a mouse inner medullary collecting duct (mIMCD3) cell line caused defective ciliogenesis [39], indicating a role for ALMS1 in the maintenance of ciliary function. In contrast, cilia appeared to assemble normally in renal collecting tubules from *ALMS1* transgenic mice [13]. Similarly, fibroblasts from human Alström syndrome patients were shown to have normal cilia but still had defects in membrane trafficking [40], indicating that the pathology observed in these patients may be due to defects in intracellular trafficking. While the role of ALMS1 in ciliary function is debatable, our previous data suggested that *ALMS1* KO rats seem to have normal ciliary development in the renal tubules indicating that the renal phenotype observed in *ALMS1* KO may be independent of cilia [11]. However, it must be noted that we only measured cilia length but not ciliary-dependent signaling in *ALMS1* KO rats [11]. Despite cilia appearing normal in size and number, vascular ciliary function such as flow-induced nitric oxide synthesis [13] may be affected in *ALMS1* KO rats, thereby causing defective vasorelaxation, as observed in this study. More studies will be needed to decipher the molecular mechanism through which ALMS1 regulates endothelial nitric oxide synthase and nitric oxide signaling in arterial smooth muscles.

We previously showed the function of ALMS1 in regulating renal function [11,41]. However, very little is known about the role of ALMS1 in other organs. ALMS1 seems to be involved in the development of obesity, since Alström syndrome patients develop early-onset obesity, and ALMS1 targeted mutations in mice lead to the development of age-dependent obesity, insulin resistance, diabetes, and hepatic steatosis between the ages of 18 to 21 weeks [13]. In foz/foz, a mouse strain carrying mutation in *ALMS1*, it was found that along with hyperphagia, impaired brown adipose tissue (BAT) diet-induced thermogenesis was evident, which is an essential factor driving diet-induced obesity and glucose intolerance [42]. Additionally, stable knockdown of ALMS1 in 3T3-L1 preadipocytes was associated with impairment of lipid accumulation and a twofold reduction in adipocyte gene expression following hormonal induction of adipogenesis [43], suggesting that partial impairment of early phase adipogenesis in Alström syndrome may contribute to the severity of the associated metabolic phenotype [43,44]. In contrast, pre-adipocytes isolated from ALMS1 mutant mice demonstrated normal adipogenic differentiation but gave rise to mature adipocytes with reduced insulin-stimulated glucose uptake. ALMS1 mutant mice also displayed altered intracellular localization of GLUT4 [45]. Reduced insulin-dependent GLUT4 translocation to the plasma membrane in adipose depots may explain glucose intolerance and compensatory hyperinsulinemia in Alström patients and in transgenic *ALMS1* mouse models [13]. Interestingly, caloric restriction in an Alström patient prevented hyperinsulinemia [46], indicating hyperinsulinemia may develop secondary to obesity, as also suggested by our data in *ALMS1* KO rats. Therefore, these observations support the idea that adipose tissue with a reduced glucose uptake may expand by de novo lipogenesis to an obese state [45].

Sex hormones are important regulators of plasma lipid metabolism. They are responsible for sexual dimorphism in the plasma lipid profile and, thus can likely account for at least part of the cardiovascular protective effect in females [1-3,16,17]. Several genes have been mechanistically linked to sex-based differences in the development of metabolic syndrome [1-3,16,17]. Some sex-based differences in the development of metabolic disorders were also seen in a mouse model of *ALMS1* mutation [13]. In this mouse model, contradictory to our data, only males became hyperglycemic. [13]. In this study, however, loss of ALMS1 in mice was not characterized for sex-based phenotypic differences in hypertension nor vascular function [13]. We previously showed that ALMS1 KO rats have elevated blood pressure and this preceded other characteristics of metabolic dysfunction in these rats [11]. Here, we show that female *ALMS1* KO rats exhibited a more severe form of metabolic syndrome than their male counterparts. The underlying cause of sex differences is unknown. However, we previously found that the carboxyl terminus of ALMS1 interacted with proteins involved in estrogen receptor signaling such as prohibitin and integrin linked kinase (ILK) [47,48], suggesting the involvement of ALMS1 in estrogen signaling. Further studies are needed to test this potential hypothesis.

## Methods

### Generation of ALMS1 KO rats

Frameshift deletion of 17 bp in exon 1 (containing a Nci1 restriction site) of ALMS1 gene was facilitated by injecting zinc Finger nucleases targeting the sequence CCCGCCTCCGACTCCGCCtccgtcCTCCCGGCACCAGTA into Dahl SS/JrHsdMcwi rat embryos. This deletion caused a frameshift leading to pre-mature stop codon in exon 2. Confirmation of deletion was done by PCR using exon 1 specific primer followed by Nci1 restriction digestion.

Forward primer-AGAGGAAGAGTTGGAAGGGG,

Reverse primer-ATACATAGGCAGAGCGACCC.

*ALMS1* KO rats were generated in Dahl salt-sensitive background and were males and females between the ages of 6-18 weeks of age. Age-matched littermate controls were male and female Dahl salt-sensitive rats, respectively. Animals were fed a standard diet (0.22% Na, 1% K) from Envigo (Indianapolis, IN). All procedures involving live animals were approved by the Institutional Animal Care and Use Committee (IACUC) of the Henry Ford Hospital and conducted following guidelines.

### Blood pressure measurements

Systolic blood pressure was also measured in awake rats by tail-cuff plethysmography.

### Plasma insulin and leptin measurement

Whole blood was collected from the abdominal aorta of rats anesthetized with Pentobarbital (Nembutal) at a dose of 100µl/100 g body weight and was immediately mixed with an appropriate volume of EDTA (5:1 ratio) to prevent blood coagulation. Blood was then spun at 200g for 20 minutes to collect plasma. Using an enzyme immunoassay kit from Cayman (Ann Arbor, MI), plasma samples were used to measure insulin and leptin concentration. Plasma renin activity Plasma renin activity was analyzed by the generation of angiotensin (Ang) I (ng Ang 88 I/ml/hr) using a Gamma Coat RIA kit (DiaSorin, Stillwater, MN) as previously described and according to the manufacturer’s instructions [49].

### Blood glucose and serum lipid profiling

Random and fasting blood glucose was measured using Wavesense Presto Blood Glucose meter (AgaMatrix, Salem, NH). Lipid profiling using rat serum samples was done using Siemen’s healthcare diagnostic analyzer (Siemens, Malvern, PA) according to the manufacturer’s instructions.

### Vascular reactivity

Briefly, rats were anesthetized with pentobarbital (50 mg/kg). Abdominal and thoracic aorta from rats was removed while maintaining endothelium intact and suspended in water baths containing perfusion solution to assess vascular reactivity. Metacholine and Nitroprusside-induced vasorelaxation was measured as a percentage of 0.1 mM Phenylephrine-induced vascular contraction.

### Statistical analysis

Results are expressed as mean ± SEM. Single intergroup comparisons between two groups were performed with a Student’s *t*-test using GraphPad Prism software. p<0.05 was considered statistically significant.

### Inclusion and Exclusion criteria

No outliers were excluded from the study. Blinding: In vascular reactivity experiments, the experimenter was blinded to the identities of rats. Power analysis: Based on previous experience, animal cohorts of 3-6 rats per group are deemed sufficient to detect differences between groups with a 90% power and a 5% type I error rate.

### Data sharing

We will follow all NIH policies concerning sharing reagents, materials, and information with other investigators. Detailed protocols are provided to everyone who requests them. Upon publication, this manuscript will be submitted to the National Library of Medicine’s PubMed Central as outlined by NIH policy.

## Acknowledgements

The authors thank the members of Cobb lab and other contributing labs for valuable suggestions, and Dionne Ware for administrative assistance. We thank Emily Henson for technical aid with blood pressure measurements. These studies were supported in part by NIH R01DK131114A1 to **PAO**, American Heart Association predoctoral fellowship 16PRE27510032 to ABJ.

## Author contribution

**ABJ**: Conceptualized, supervised, designed and performed experiments, performed analysis, wrote the manuscript, generated initial figures; **SRM**: Collected and analyzed data; **NER**: performed experiments and edited the manuscript; **MM**: performed experiments; **PAO**: Supervision, acquired funding, edited manuscript.

## Declaration of interests

The authors declare no competing interests.

## Notes

### Competing Interest Statement

The authors have declared no competing interest.

## References

1) Ryan AS. Insulin resistance with aging: effects of diet and exercise. Sports Med. 2000 Nov;30(5):327–46.

2) American Diabetes Association. 12. Older Adults: Standards of Medical Care in Diabetes-2020. Diabetes Care. 2020 Jan;43(Suppl 1):S152–S162.

3) Geer EB, Shen W. Gender differences in insulin resistance, body composition, and energy balance. Gend Med. 2009;6 Suppl 1(Suppl 1):60–75.

4) Marshall JD, Hinmam EG, Collin GB, Beck S, Cerqueira R, Maffei P, Milan G, Zhang W, Wilson DI, Hearn T, Tavares P, Vettor R, […], So WV, Nishina PM, Naggert JK. Spectrum of ALMS1 variants and evaluation of genotype-phenotype correlations in Alström syndrome. Hum Mutat. 28:1114–1123, 2007.

5) Marshall JD, Bronson RT, Collin GB, Nordstrom AD, Maffei P, Paisey RB, Carey C, MacDermott S, Russell-Eggitt I, Shea SE, Davis J, Beck S, Sicolo N, Naggert JK, Nishina PM. New Aström Syndrome phenotypes based on the evaluation of 182 cases. Arch Intern Med. 165:675–683, 2005.

6) Marshall JD, Maffei P, Collin GB, Naggert JK. Alström syndrome: Genetics and clinical overview. Curr Genomics. 12:225–235, 2011.

7) Joy T, Cao H, Black G, Malik R, Charlton-Menys V, Hegele RA, Durrington PN. Alström syndrome (OMIM 203800): a case report and literature review. Orphanet J Rare Diseases. 2:49, 2007.

8) Go AS, Chertow GM, Fan d, McCulloch CE, Hsu CY. Chronic kidney disease and the risks of death, cardiovascular events and hospitalization. N Engl J Med. 351:1296–305, 2004.

9) Kastarinen M, Juutilainen A, Kastarinen H, Salomaa V, Karhapaa P. Risk factors for end-stage renal disease in a community-based population: 26-year follow-up of 25,821 men and women in eastern Finland. J Intern Med. 267:612–20, 2010.

10) Barkley RA, Chakravarti A, Cooper RS, Ellison C, Hunt SC, Province MA, Turner ST, Weder AB, Boerwinkle E. Positional Identification of hypertension 104 susceptibility genes on chromosome 2. Hypertension. 43:477–482, 2004.

11) Jaykumar AB, Caceres PS, King-Medina KN, Liao TD, Datta I, Maskey D, Naggert JK, Mendez M, Beierwaltes WH, Ortiz PA. Role of Alström syndrome 1 in the regulation of blood pressure and renal function. JCI Insight. 2018 Nov 2;3(21):e95076.

12) Favaretto F, Milan G, Collin GB, Marshall JD, Stasi F, Maffei P, Vettor R, Naggert JK. GLUT4 defects in adipose tissue are early signs of metabolic altercations in Alms1GT/GT, a mouse model for obesity and insulin resistance. PLoS One. 9:e109540, 2014.

13) Collin GB, Cyr E, Bronson R, Marshall JD, Gifford EJ, Hicks W, Murray SA, Zheng QY, Smith RS, Nishina PM, Naggert JK. Alms1-disrupted mice recapitulate human Alström syndrome. Hum Mol Genet. 2005 Aug 15;14(16):2323–33.

14) Yoon SS, Carroll MD, Fryar CD. Hypertension prevalence and control among adults: United States, 2011-2014. NCHS Data Brief. 220:1–8, 2015.

15) Heidenreich PA, Trogdon JG, Khavjou OA, Butler J, Dracup K, Ezekowitz MD, Finkelstein EA, Hong Y, Johnston SC, Khera A et al. Forecasting the future of cardiovascular disease in the united states: a policy statement from the American Heart Association. Circulation. 123:933–944, 2011.

16) Kannel WB. McGee DL. Diabetes and cardiovascular risk factors: the Framingham study. Circulation. 59:8–13, 1979.

17) Stamler J, Vaccaro O, Neaton JD, Wentworth D. Diabetes, other risk factors and 12-yr cardiovascular mortality for men screened in the Multiple Risk Factor Intervention Trial. Diabetes Care 16:434–444, 1993.

18) Frohlich ED. Clinical management of the obese hypertensive patient. Cardiol Rev. 10(3):127–138, 2002.

19) Hall JE. Pathophysiology of obesity hypertension. Curr Hypertens Rep. 2(2):139–147, 2000.

20) Reckelhoff JF. Gender differences in the regulation of blood pressure. Hypertension. 2001 May;37(5):1199–208.

21) Hall JE, Crook ED, Jones DW, Wofford MR, Dubbert PM. Mechanisms of obesity-associated cardiovascular and renal disease. Am J Med Sci. 324(3):127–137, 2002.

22) Albillos A, Rossi I, Cacho G, Martínez MV, Millán I, Abreu L, Barrios C, Escartín P. Enhanced endothelium-dependent vasodilation in patients with cirrhosis. Am J Physiol. 1995 Mar;268(3 Pt 1):G459–64.

23) Frohlich ED. Clinical management of the obese hypertensive patient. Cardiol Rev. 10(3):127–138, 2002.

24) Hall JE. Pathophysiology of obesity hypertension. Curr Hypertens Rep. 2(2):139–147, 2000.

25) Hall JE, Crook ED, Jones DW, Wofford MR, Dubbert PM. Mechanisms of obesity-associated cardiovascular and renal disease. Am J Med Sci. 324(3):127–137, 2002.

26) Wofford MR, Hall JE. Pathophysiology and treatment of obesity hypertension. Curr Pharm Des. 10(29):3621–3637, 2004. 116

27) Bakris GL, Weir MR, Sowers JR. Therapeutic challenges in the obese diabetic patient with hypertension. Am J Med. 101:33S–46S, 1996.

28) Köttgen A, Pattaro C, Boger CA, Fuchsberger C, Olden M, Glazer NL, Parsa A, Gao X, Yang Q, Smith AV, O’Connell JR, Li M, […], Kao WH, Heid IM and Fox CS. Multiple new loci associated with kidney function and chronic kidney disease: The CKDGen consortium. Nature Genet. 42:376–384, 2010.

29) Chambers JC, Zhang W, Lord GM. Genetic loci influencing kidney function and chronic kidney disease. Nature Genetics. 42:373–75, 2010. 105

30) Boger CA, Heid IM. Chronic kidney disease: novel insights from genome-wide association studies. Kidney Blood Press Res. 34:225–234, 2011.

31) Boger CA et al. Association of eGFR-related loci identified by GWAS with incident CKD and ESRD. PLoS Genet. 7(9):e1002292, 2011.

32) Schulz A, Litfin A, Kossmehl P, Kreutz R. Genetic dissection of increased urinary albumin excretion in the munich wistar fromter rat. J Am Soc Nephrol. 13:2706–2714, 2002.

33) Atwood LD, Samollow PB, Hixson JE, Stern MP, MacCluer JW. Genome-wide linkage analysis of blood pressure in Mexican Americans. Genet. Epidemiol. 20:373–382, 2001.

34) Aneas I, Rodrigues MV, Pauletti BA, Silva GJJ, Carmona R, Cardoso L, Kwitek AE, Jacob HJ, Soler JMP, Krieger JE. Congenic strains provide evidence that four mapped loci in chromosomes 2, 4, and 16 influence hypertension in the SHR. Physiol Genomics. 37:52–57, 2009.

35) Sugiyama F, Churchill GA, Higgins DC, Johns C, Makaritsis KP, Gavras H, Paigen B. Concordance of murine quantitative trait loci for salt-induced hypertension with rat and human loci. Genomics. 71:70–77, 2001.

36) Takami S, Higaki J, Rakugi H, Miki T, Katsuya T, Nakata Y, Serikawa T, Ogihara T. Analysis and comparison of new candidate loci for hypertension between genetic hypertensive rat strains. Hypertens Res. 19:51–56, 1996.

37) McKay EJ, Luijten I, Weng X, Martinez de Morentin PB, De Frutos González E, Gao Z, Kolonin MG, Heisler LK, Semple RK. Mesenchymal-specific Alms1 knockout in mice recapitulates metabolic features of Alström syndrome. Mol Metab. 2024 Jun;84:101933.

38) Heydet D, Chen LX, Larter CZ, Inglis C, Silverman MA, Farrell GC, Leroux MR. A truncating mutation of Alms1 reduces the number of hypothalamic neuronal cilia in obese mice. Dev Neurobiol. 73(1):1–13, 2013.

39) Li G, Vega R, Nelms K, Gekakis N, Goodnow C, McNamara P, Wu H, Hong NA, Glynne R. A role for Alström syndrome protein, Alms1, in kidney ciliogenesis and cellular quiescence. PLoS Genet. 3(1):e8, 2007.

40) Collin GB, Marshall JD, King BL, Milan G, Maffei P, Jagger DJ, Naggert JK. The Alström syndrome protein, ALMS1, interacts with α-Actinin and components of the endosome recycling pathway. PLoS One. 7:e37925, 2012.

41) Monu SR, Potter DL, Liao TD, King KN, Ortiz PA. Role of Alström syndrome 1 in the regulation of glomerular hemodynamics. Am J Physiol Renal Physiol. 2023 Oct 1;325(4):F418–F425.

42) Festuccia WT. Turning up the heat against metabolic syndrome and nonalcoholic fatty liver disease. Clin Sci (Lond). 131(4):327–328, 2017.

43) Huang-Doran I, Semple RK. Knockdown of Alström syndrome-associated gene Alms1 in 3T3-L1 predipocytes impairs adipodenesis but has no effect on cell autonomous insulin action. Int J Obes (Lond). 34(10):1554–8, 2010.

44) Romano S, Milan G, Veronese C, Collin GB, Marshall JD, Centobene C, Favaretto F, Dal Pra C, Scarda A, Leandri S, Naggert JK, Maffei P, Vettor R. Regulation of Alström syndrome gene expression during adipogenesis and its relationship with fat cell insulin sensitivity. Int J Mol Med. 21(6):731–736, 2008. 115

45) Favaretto F, Milan G, Collin GB, Marshall JD, Stasi F, Maffei P, Vettor R, Naggert JK. GLUT4 defects in adipose tissue are early signs of metabolic altercations in Alms1GT/GT, a mouse model for obesity and insulin resistance. PLoS One. 9:e109540, 2014.

46) Lee NC, Marshall JD, Collin GB, Naggert JK, Chien YH, Tsai WY, Hwu WL. Caloric restriction in Alström syndrome prevents hyperinsulinemia. Am J Med Genet A. 149A(4):666, 2009.

47) He B, Feng Q, Mukherjee A, Lonard DM, DeMayo FJ, Katzenellenbogen BS, Lydon JP, O’Malley BW. A repressive role for prohibitin in estrogen signaling. Mol Endocrinol. 2008 Feb;22(2):344–60.

48) Acconcia F, Manavathi B, Mascarenhas J, Talukder AH, Mills G, Kumar R. An inherent role of integrin-linked kinase-estrogen receptor alpha interaction in cell migration. Cancer Res. 2006 Nov 15;66(22):11030–8.

49) Mendez M, Gaisano HY. Role of the SNARE protein SNAP23 on cAMP-stimulated renin release in mouse juxtaglomerular cells. Am J Physiol Renal Physiol. 2013 Mar 1;304(5):F498–504.

